# The genome of the rayed Mediterranean limpet *Patella caerulea* (Linnaeus, 1758)

**DOI:** 10.1101/2023.12.18.572124

**Authors:** Gwyneth Halstead-Nussloch, Silvia Giorgia Signorini, Marco Giulio, Fabio Crocetta, Marco Munari, Camilla Della Torre, Alexandra Anh-Thu Weber

## Abstract

*Patella caerulea* (Linnaeus, 1758) is a molluscan limpet species of the class Gastropoda. Endemic to the Mediterranean Sea, it is considered to be a keystone species in tidal and subtidal habitats due to its primary role in structuring and regulating the ecological balance of these habitats. It is currently being used as a bioindicator to assess the environmental quality of coastal marine waters and as a model species to understand adaptation to ocean acidification. Here we provide a high-quality reference genome assembly and annotation for *Patella caerulea*. We used a single specimen collected in the field to generate ∼30 Gb of PacBio HiFi data. The final assembly is 749.8 Mb large and contains 62 contigs, including the mitochondrial genome (14,938 bp). With an N50 of 48.8 Mb and 98% of the assembly contained in the 18 largest contigs, this assembly is near chromosome-scale. BUSCO scores were high (Mollusca: 87.8% complete; Metazoa: 97.2% complete) and similar to metrics observed for other chromosome-level *Patella* genomes, highlighting a possible bias in the Mollusca database for Patellids. We generated transcriptomic Illumina data from a second individual collected at the same locality, and used it together with protein evidence to annotate the genome. 23,938 protein coding gene models were found. By comparing this annotation with other published *Patella* annotations, we found that the distribution and median values of exon and gene lengths was comparable to other *Patella* species despite different annotation approaches. The present high-quality *Patella caerulea* reference genome is an important resource for future ecological and evolutionary studies.

**Significance:** Reference genomes are essential resources for biodiversity conservation and management. *Patella caerulea* (Linnaeus, 1758) is a gastropod species occurring in the Mediterranean Sea that is currently used as a model to understand the impact of pollution and ocean acidification on marine biodiversity. Here we present a high-quality reference genome of *P. caerulea*, that almost reaches chromosome-level contiguity. We further provide a high-quality genome annotation supported by transcriptomic evidence. This reference genome will be of interest for researchers working on the ecology and evolution of marine biodiversity. All data is available on the public database NCBI for future use by researchers.

## Introduction

Patellid limpets (family Patellidae Rafinesque, 1815) are gastropod molluscs, containing approximately 400 species distributed worldwide. Patellid species are divided into four different genera (Ridgway et al., 1998; Koufopanou et al., 1999). Species of the genus *Patella* mostly occur in the north-eastern Atlantic and the Mediterranean Sea, and this genus currently includes 16 valid species. Some of these species have a very limited distribution and are endemic to isolated islands and archipelagos (Titselaar, 2019), while others have a wider distribution, being widespread in the Atlantic Ocean and/or the Mediterranean Sea (Gofas et al., 2012; Alf et al., 2020). However, the correct taxonomy of some taxa is still under investigation, as they may account for yet-unsolved species complexes (Sá-Pinto et al., 2010). The rayed Mediterranean limpet *Patella caerulea* (Linnaeus, 1758) is endemic to the Mediterranean Sea and widespread from the western to the eastern shores of the basin (Grossu, 1993; Gofas et al., 2012; Crocetta et al., 2020). Shell morphological traits of this species are highly variable, with the presence of different morphs depending on ontogenic or ecophenotypic variables (Cossignani and Ardovini, 2011; Gofas et al., 2012). This is reflected in a long list of synonyms and even varieties described by past authors, now all synonymized under the valid nominal taxon (MolluscaBase eds., 2023). Notwithstanding such limitations, *P. caerulea* is generally characterized by an external greyish-brownish shell, with darker radiating bands and thin radial costae, and by the presence of a bluish iridescent layer internally, often well evident and from which the species is named. The species is protandrous hermaphrodite and is considered a predominantly winter breeder (Ferranti et al., 2018). *Patella caerulea* lives attached to rocks, from the lower intertidal to the subtidal zone, usually up to 5 meters depth. It is considered a keystone species due to its primary role in structuring and regulating the ecological balance of intertidal communities, both directly through grazing and acting as prey for higher trophic-level consumers, and indirectly by enhancing or inhibiting the settlement of other organisms (Menge, 2000; Reguera et al., 2018; Vafidis et al., 2020; AYDIN et al., 2021). Similar to other limpets, the species is threatened by different environmental and anthropogenic disturbances, which include ocean acidification, rising temperature, and pollution (Vafidis et al., 2020).

Given the wide distribution and abundance, sedentary lifestyle, and key ecological role, *P. caerulea* has been used as bioindicator in studies aimed at assessing the environmental quality of coastal marine waters, through the measurement of bioaccumulation in tissues and the adverse biological effects of different classes of xenobiotics (Aydin-Önen and Öztürk, 2017; Reguera et al., 2018; Zaidi et al., 2022). In this context, a study investigated the potential influence of coastal urbanization on the genetic variation of *P. caerulea* (Fauvelot et al., 2009). This study highlighted the fact that coastal urbanization may act as a biotic homogenization, reducing genetic diversity of local species, with consequences on productivity, growth, stability and interactions at both community and ecosystem levels. Finally, *P. caerulea* is currently used as a model to understand adaptation to ocean acidification in gastropods, as natural populations adapted to reduced pH conditions (∼7.4) have been discovered in the CO_2_ vent systems of Ischia Island, Italy together with *P. ulyssiponensis* and *P. rustica* (Aliende et al., 2023). Specifically, individuals of these three species occurring in reduced pH conditions have thinner shells and larger sizes than individuals from ambient pH conditions. However, the specific molecular mechanisms underlying this phenotypic variability are unknown. Specifically, the respective influence of phenotypic plasticity vs. genetic adaptation has not been examined yet, and is currently under investigation (Signorini et al, in preparation).

Despite its important ecological and evolutionary roles and its use as a bioindicator, there is to date no reference genome for *P. caerulea*. There are currently four reference genome projects of *Patella* species, three that are published and assembled to the chromosome level (*P. depressa, P. vulgata, P. pellucida*) (Lawniczak et al., 2022; Hawkins et al., 2023) and one (*P. ulyssiponensis)*, led by the Darwin Tree of Life initiative, for which the chromosome-scale genome assembly has been recently released (ENA project PRJEB63446), but the genome note has to date not been published (Table 1). These *Patella* species are diploid, have between eight and nine pairs of chromosomes (Petraccioli et al., 2010), and have assembly sizes varying between 683 Mb and 712 Mb. *P. caerulea* also has nine pairs of chromosomes (Petraccioli et al., 2010), but there are no genome size estimates for this species.

**Table 1:**
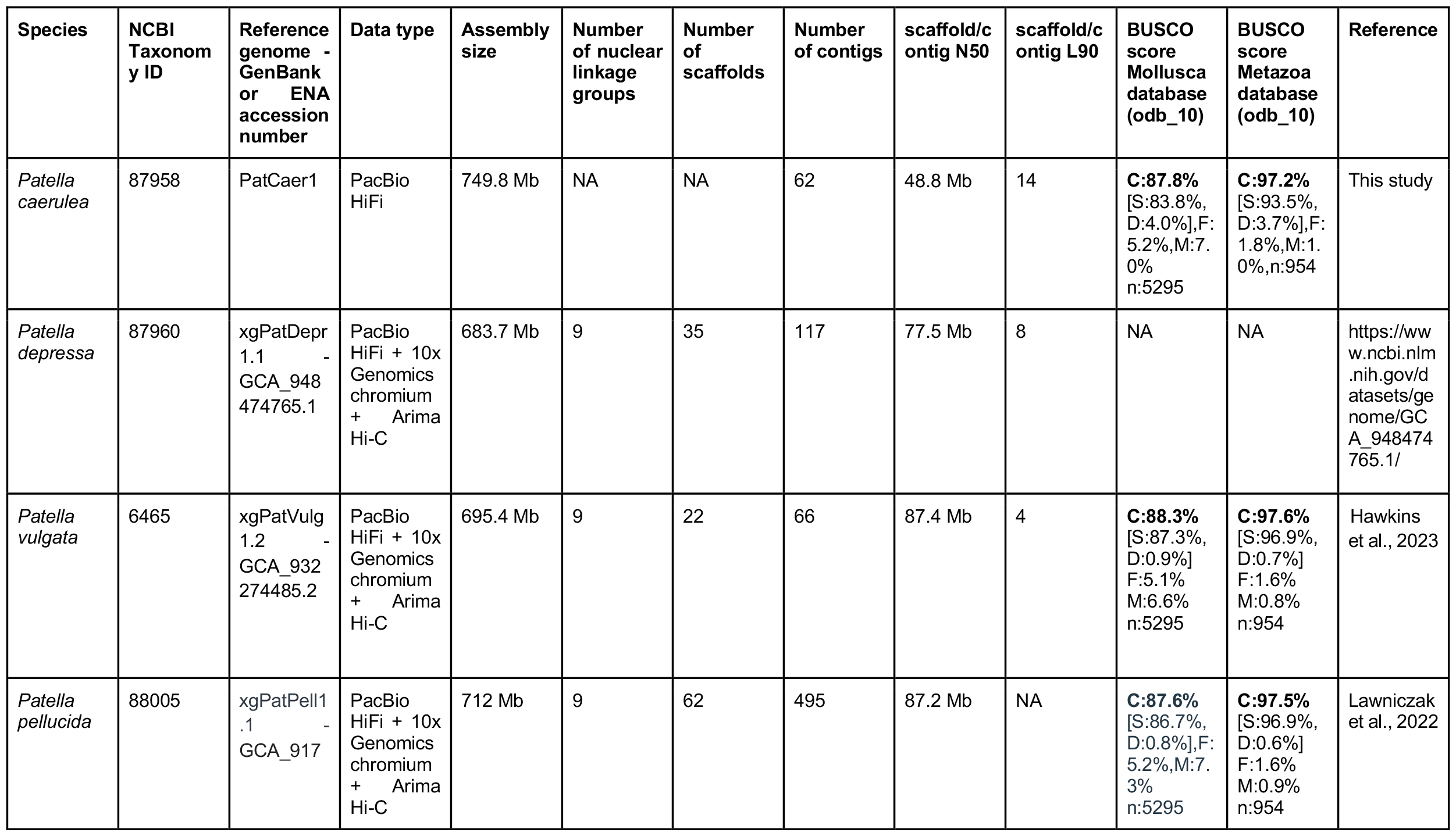
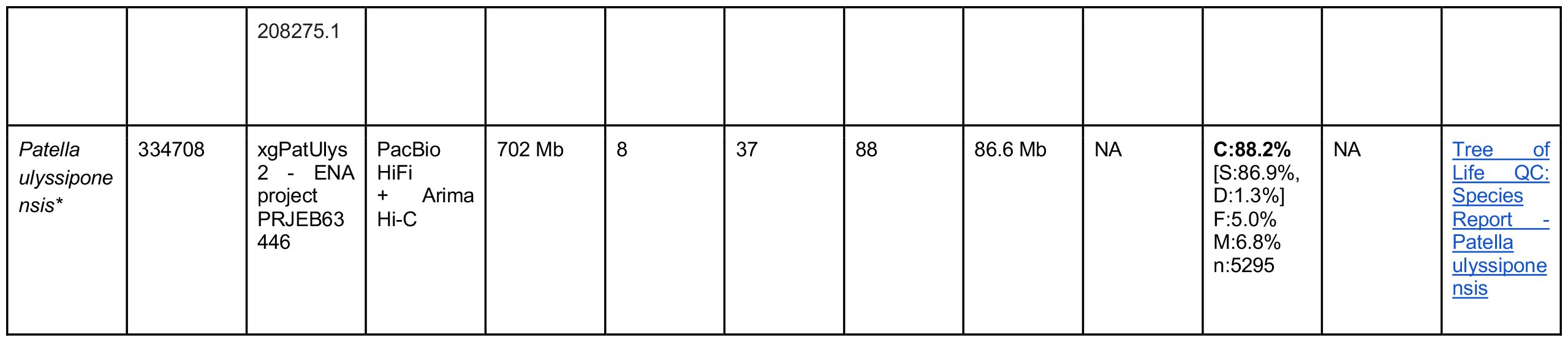
Metrics of the *Patella caerulae* PatCaer1 genome assembly and comparison with publicly available *Patella* genome projects. *Assembly statistics may change as the *P. ulyssiponensis* genome note is unpublished yet (status 08.12.2023).

Having high-quality reference genomes is essential for the broad field of biodiversity genomics (Formenti et al., 2022; Theissinger et al., 2023). As a result, numerous initiatives and consortia are now aiming at generating high-quality reference genomes for all described eukaryotic species on Earth (e.g. the Earth Biogenome Project - EBP) (Lewin et al., 2018, 2022). The European Reference Genome Altas (ERGA) initiative is the European node of the EBP, which aims at sequencing and assembling the genomes of all eukaryotic species in Europe (Mazzoni et al., 2023; Mc Cartney et al., 2023). Within animals, while sequencing of vertebrates is advancing well (Hotaling et al., 2021; Rhie et al., 2021), genome projects on invertebrates, especially arthropods and molluscs, are comparatively lagging behind (Hotaling et al., 2021). Here we provide a high-quality reference genome of *Patella caerulea* which will help future ecological and evolutionary studies of the rayed Mediterranean limpet.

## Results and Discussion

### Genome assembly

We used a single *P. caerulea* specimen to generate about 30 Gb of HiFi PacBio data from a single SMRT cell. Genome size was estimated to be ∼623 Mb and heterozygosity of 2.06% using k-mer based methods (Fig. 1A). Correctness (QV) was estimated to be 68.5 and k-mer completeness was 77.53% (Fig. 1B), similar to haploid assemblies of other heterozygous species (Qi et al., 2022). The final assembly was 749.8 Mb in length and contained 62 contigs including the mitochondrial genome (Table 1, Fig. 1C). Assuming that the genome size is closer to the assembly size than the genome scope estimate, this results in an approximate sequencing coverage of 40x. 98.4% of the assembly was contained in the 18 largest contigs (Fig. 1D), indicating an assembly close to chromosome-level from HiFi reads alone, given that *P. caerulea* has nine pairs of chromosomes. BUSCO completeness was 87.8% using the mollusca_odb10 and 97.2% with metazoa_odb10 reference set, a pattern also observed for at least three other *Patella* species (Table 1), further indicating potential clade specific biases in the mollusca reference set. This may indicate lineage-specific gene losses, which could also be the case for other species of the Patellidae family in future reference genome projects. These metrics show that this very contiguous assembly provides a high-quality genomic resource for *Patella caerulea* generated with data from a single sequencing technology (PacBio). A 14,938 bp contig corresponding to the mitochondrial genome was identified within the primary assembly by Tiara classification and confirmed using BLAST and presence of mitochondrial genes using MITOS2 (Donath et al., 2019). Another contig identified as mitochondrial by Tiara had subsequent BLAST hits with other *Patella* nuclear (i.e. ribosomal genes) but not mitochondrial genome sequences. We therefore kept this contig in the nuclear assembly.

**Figure 1.**
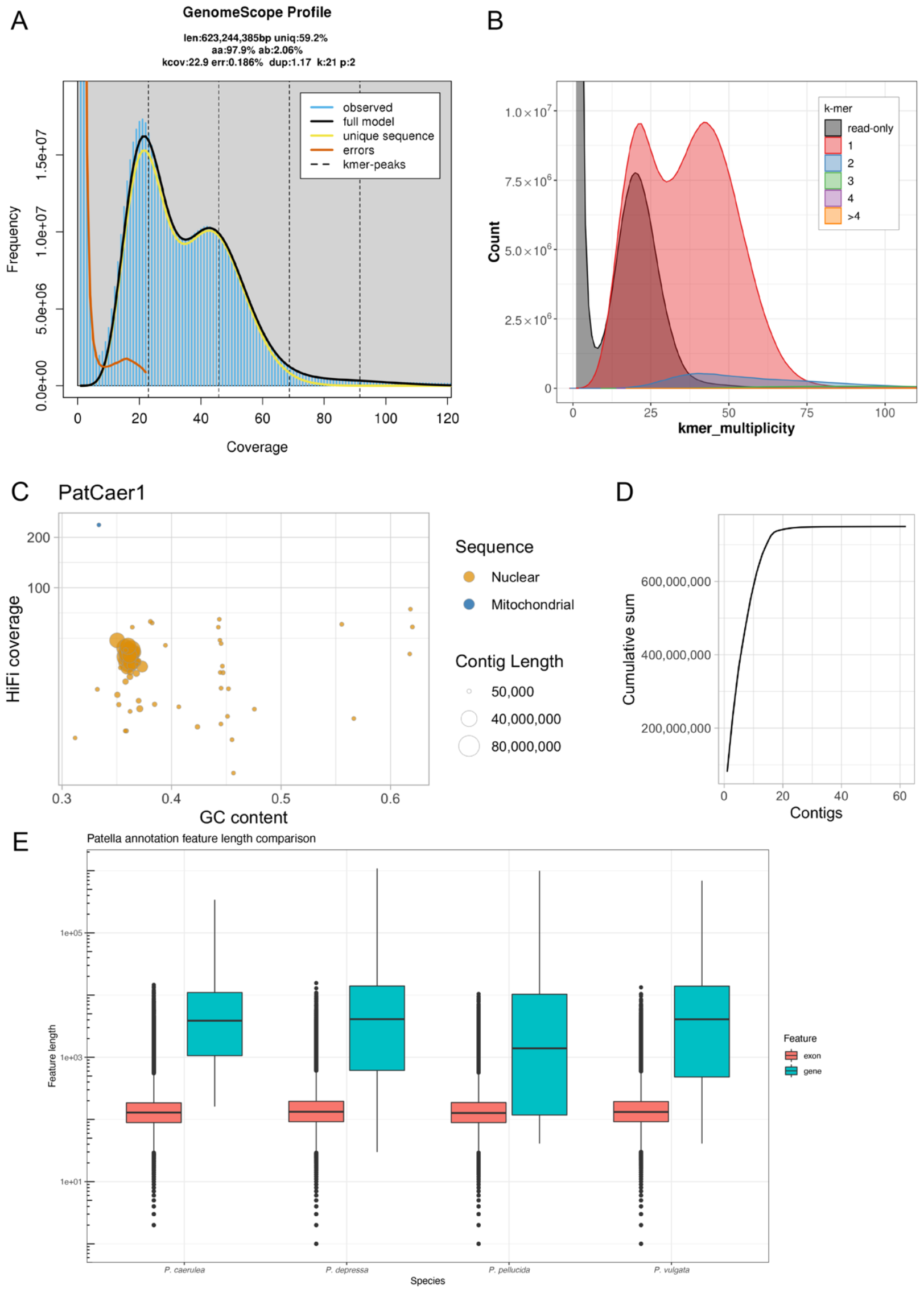
*Patella caerulea* PatCaer1 genome assembly and annotation metrics. **A)** GenomeScope2 k-mer profile with estimates of genome size and heterozygosity. **B)** Merqury k-mer plot. Assembly k-mer copy number spectrum shows low level of duplications within the assembly and that most kmers are incorporated into the haploid assembly. Read-only k-mers in the single-copy peak are from alternate haplotypes from this heterozygous species. **C)** Contigs were evaluated by mapping HiFi reads back on the assembly and calculating mean coverage for each contig (y-axis: HiFi coverage). Contig average GC content (x-axis), Tiara sequence classification (circle color) and length (circle size) is also displayed. **D)** Cumulative sum of contig length (bp), ordered by contig length. **E)** Annotation feature comparison with other *Patella* annotations available on NCBI. Distribution and median values for gene and exon length are similar between species.

### Genome annotation

Gene models were generated using both RNA and protein evidence, which resulted in 23,938 protein coding gene models (Table S1). This number is higher than the number of protein coding genes identified in *Patella vulgata*, where the most similar annotation pipeline was implemented (i.e. the use of BRAKER2) (Hawkins et al., 2023). BUSCO completeness was 90.7% using the mollusca_odb10 database and 96.0% with metazoa_odb10, similar or even higher than the completeness assessed for the genome assembly, which suggests high-quality annotation (Manni et al., 2021b). The distribution and median values of exon and gene lengths was comparable to other *Patella* species despite different annotation approaches, indicating congruence in structural feature identification within the *Patella* assemblies (Fig. 1E). It should be noted that annotations for the other *Patella* species include further classifications beyond protein coding sequences that were not identified in this version of the *P. caerulea* annotation, increasing the number of gene and transcript features in those annotations (Table S1). Finally, we found 16,646 reciprocal best blast hits between *P. caerulea* and *P. depressa* amino acid sequences.

We note that while genome assembly metrics are easily comparable between projects and datasets, it is more difficult to compare genome annotations, even between species from the same genus. While there are initiatives to standardize annotations (e.g. ERGA recommendations https://www.erga-biodiversity.eu/structural-annotation), it is currently difficult to directly compare annotations between species, especially non-model organisms. Indeed, annotation pipelines utilize different methods and availability (or lack thereof) of external evidence (e.g. RNAseq), which quickly generates discrepancies between each annotation project. Finally, we acknowledge that Hi-C data would add valuable information to further scaffold to the chromosome level and phase the present assembly. It would as well be useful for manual curation and correction of potential misassemblies. However, the present primary assembly is already highly contiguous and well-annotated, and we believe it will be a valuable resource for the community of researchers working on the ecology and evolution of *Patella caerulea*.

## Materials and Methods

### Sample collection, DNA and RNA extractions, sequencing

Ten *Patella caerulea* individuals were collected by snorkeling at 2m depth on 9th of June 2023 in Punta San Pietro, Ischia Island, Italy [40.74702561880045, 13.943306501480043] with the permission of the Marine Protected Area ‘Regno di Nettuno’. High-molecular weight DNA extraction from a single individual (NCBI Biosample SAMN38441487, PatCaer1) was performed using the Qiagen MagAttract HMW kit following manufacturer’s instructions. The individual was not sexed. A single PacBio standard library (15-20 kb insert size) was constructed and sequenced on one SMRT cell 8M on a PacBio Sequel II instrument at the Functional Genomics Center Zürich (FGCZ), Switzerland. A second specimen (NCBI Biosample SAMN38441544, PatCaer2) was used for RNA extractions. Individual RNA extractions from head, foot/mantle and visceral body were performed using the E.Z.N.A.® HP Total RNA Isolation Kit (Omega Bio-tek). Three RNA extractions passing QC (RIN > 9) were pooled in equimolar concentrations and a single transcriptomic library was constructed using Truseq mRNA kit (Illumina) and sequenced on a Illumina Novaseq 6000 sequencer (200M reads, 150bp paired-end) at the FGCZ.

### Genome size estimate and genome assembly

1,984,661 Pacific Biosciences high fidelity (PacBio HiFi) reads containing a total of 29,667,167,600 bases and a mean length of 14,948 bp were used for the assembly. Meryl v1.3 (Rhie et al., 2020) was used to generate a k-mer database. K-mer frequency was analyzed and genome size estimated by GenomeScope2 (Ranallo-Benavidez et al., 2020). A primary assembly was produced using hifiasm (Cheng et al., 2022) v0.19.6-r595 with purging parameter set to -s 0.35. Contigs were evaluated by: 1) measuring GC content using asmstats (https://github.com/marcelauliano/Teaching/blob/main/asmstats) and seqtk v1.3-r117-dirty (https://github.com/lh3/seqtk), 2) coverage of HiFi reads mapped back to the assembly using minimap2 (Li, 2018), and 3) sequence classification by Tiara (Karlicki et al., 2021). Contigs not classified as eukaryotic or mitochondrial, with coverage below 5x or above 100x were removed from the primary assembly. Assembly metrics were further evaluated using BUSCO v5.2.2 with both the mollusca and metazoa odb10 databases (Manni et al., 2021a). QV, k-mer completeness and k-mer copy number plots were estimated and generated using merqury v1.3 (Rhie et al., 2020).

### Annotation

Prior to annotation, repeat libraries including LTR sequences were generated from the contigs using RepeatModeler2 (Flynn et al., 2020) and combined with mollusca specific repeat family sequences from DFam (Storer et al., 2021) prior to running RepeatMasker (Smit et al., 2013). The softmasked assembly was then used as input for the BRAKER pipelines (Hoff et al., 2016, 2019; Bruna et al., 2021) in order to generate gene models. Briefly, 200M paired-end RNAseq reads were aligned to the masked assembly using Hisat2 (Kim et al., 2019) and the bam file was used as input for the BRAKER1 pipeline (Hoff et al., 2016). Metazoa specific protein sequences partitioned from OrthoDB v11 (Kuznetsov et al., 2023) were used as input for the BRAKER2 pipeline (Bruna et al., 2021). The gene models in the braker.gtf output from both pipelines were combined using default evidence weights, the --ignore_tx_phase option and renamed using TSEBRA (Gabriel et al., 2021).

We followed some recommendations of the ERGA initiative to assess the quality of our annotation (https://www.erga-biodiversity.eu/structural-annotation). Specifically, we used BUSCO v5.2.2 with both the mollusca and metazoa odb10 databases (Manni et al., 2021a) to evaluate the completeness of our annotation, using the longest isoform for analysis. We also evaluated our annotation by comparing the distribution and median lengths of features classified as “exon” and “gene” among the species *P. caerulea, P. depressa, P. pellucida* and *P. vulgata* for which annotation data is available on NCBI. Finally, we ran a reciprocal best blast hit analysis between the *P. caerulea* and *P. depressa* amino acid sequences to infer how many orthologous genes are present between both species.

## Data availability

The genomic and transcriptomic raw data are available on NCBI SRA Bioproject ID PRJNA1045377 (https://www.ncbi.nlm.nih.gov/sra): Biosample IDs SAMN38441487 (for PacBio HiFi), SAMN38441544 (for RNAseq); PacBio data: SRX22853381, RNAseq data: SRX22853382. The PatCaer1 genome assembly and annotation have been deposited on NCBI (https://www.ncbi.nlm.nih.gov/genome/) and are currently being processed. The commands used are available on the GitHub repository https://github.com/GwynHN/PatCaer_asm_anno.

## Acknowledgments

We are grateful to Director Antonio Miccio and Caterina Iacono of the MPA ‘Regno di Nettuno’ of Ischia. We would like to thank Pascal Bucher for his help in the Aquatikum facility. Sequencing was performed at the Functional Genomics Center Zürich (FGCZ), Switzerland. Data produced and analyzed in this paper were generated in collaboration with the Genetic Diversity Centre (GDC), ETH Zürich.

## Funding

The project was funded by the Swiss Federal Institute of Aquatic Science and Technology (Eawag).

## Author contributions

GHN: genome assembly and annotation, manuscript drafting; SS, CDT, MM, FC: sample acquisition, manuscript drafting and editing; MG: laboratory work; AATW: supervision, funding, manuscript drafting and editing.

---

**Table S1.** Annotation metrics of *P. caerulea* and comparison with published *Patella* genome annotations. See Table 1 for references.

| Statistics | *P. caerulea* | *P. depressa* | *P. pellucida* | *P. vulgata* |
| --- | --- | --- | --- | --- |
| Number of protein-coding genes | 23,938 | 20,502 | 18,161 | 19,378 |
| Number of transcripts | 29,627 | 40,054 | 53,889 | 47,283 |
| Mean gene length | 9,284 | 13,421 | 10,190 | 12,757 |
| Median gene length | 3,870 | 4,098 | 1,393 | 4,088 |
| Mean exon length | 208 | 211 | 201 | 214 |
| Median exon length | 129 | 133 | 127 | 132 |
| Mean exons per transcript | 7.09 | 6.25 | 5.55 | 6.54 |
| Median exons per transcript | 4 | 3 | 2 | 3 |
| BUSCO score Mollusca (odb10) | C:90.7%<br>[S:86.9%,D:3.8%],<br>F:0.8%,M:8.5%,n:5295 | NA | NA | NA |
| BUSCO score Metazoa (odb10) | C:96.0%<br>[S:92.2%,D:3.8%],<br>F:1.2%,M:2.8%,n:954 | NA | NA | NA |

## Notes

https://www.ncbi.nlm.nih.gov/bioproject/?term=PRJNA1045377

### Competing Interest Statement

The authors have declared no competing interest.

## References

Alf, A., Brenzinger, B., Haszprunar, G., Schrodl, M., Schwabe, E. (2020). A guide to marine molluscs of Europe. ConchBooks Eds. 803 pp.

Aliende, M., Busselo, A., NarÉ, M., Sahuquillo, M., Iacono, G., Iacono, C., et al. (2023). ABUNDANCE AND SIZE STRUCTURE OF PA?LA SPP.(MOLLUSCA, GASTROPODA) UNDER OCEAN ACIDIFICATION CONDITIONS AT CO2 VENTS (ISCHIA ISLAND, ITALY). Biol. Mar. Mediterr. 27, 14.

Aydin, M., Şahin, A. E., and KaradurmuŞ, U. (2021). Some biological parameters of Patella caerulea from the Black Sea. Mar. Sci. Technol. Bull. 10, 396–405.

Aydin-Önen, S., and Öztürk, M. (2017). Investigation of heavy metal pollution in eastern Aegean Sea coastal waters by using Cystoseira barbata, Patella caerulea, and Liza aurata as biological indicators. Environ. Sci. Pollut. Res. 24, 7310–7334. doi: 10.1007/s11356-016-8226-4.

Bruna, T., Hoff, K. J., Lomsadze, A., Stanke, M., and Borodovsky, M. (2021). BRAKER2: automatic eukaryotic genome annotation with GeneMark-EP+ and AUGUSTUS supported by a protein database. NAR Genomics Bioinforma. 3, qaa108.

Cheng, H., Jarvis, E. D., Fedrigo, O., Koepfli, K.-P., Urban, L., Gemmell, N. J., et al. (2022). Haplotype-resolved assembly of diploid genomes without parental data. Nat. Biotechnol. 40, 1332–1335. doi: 10.1038/s41587-022-01261-x.

Cossignani, T., and Ardovini, R. (2011). Malacologia mediterranea: atlante delle conchiglie del Mediterraneo: 7500 foto a colori. L’informatore Piceno.

Crocetta F., Bitar G., Zibrowius H., Oliverio M. (2020). Increase in knowledge of the marine gastropod fauna of Lebanon since the 19th century. Bull. Mar. Sci. 96, 1–22.

Donath, A., Jühling, F., Al-Arab, M., Bernhart, S. H., Reinhardt, F., Stadler, P. F., et al. (2019). Improved annotation of protein-coding genes boundaries in metazoan mitochondrial genomes. Nucleic Acids Res. 47, 10543–10552. doi: 10.1093/nar/gkz833.

Fauvelot, C., Bertozzi, F., Costantini, F., Airoldi, L., and Abbiati, M. (2009). Lower genetic diversity in the limpet Patella caerulea on urban coastal structures compared to natural rocky habitats. Mar. Biol. 156, 2313–2323. doi: 10.1007/s00227-009-1259-1.

Ferranti, M. P., Monteggia, D., Asnaghi, V., and Chiantore, M. (2018). Artificial reproduction protocol, from spawning to metamorphosis, through noninvasive methods in Patella caerulea Linnaeus, 1758. Aquac. Res. 49, 3386–3391. doi: 10.1111/are.13802.

Flynn, J. M., Hubley, R., Goubert, C., Rosen, J., Clark, A. G., Feschotte, C., et al. (2020). RepeatModeler2 for automated genomic discovery of transposable element families. Proc. Natl. Acad. Sci. U. S. A. 117, 9451–9457. doi: 10.1073/pnas.1921046117.

Formenti, G., Theissinger, K., Fernandes, C., Bista, I., Bombarely, A., Bleidorn, C., et al. (2022). The era of reference genomes in conservation genomics. Trends Ecol. Evol. 37, 197–202. doi: 10.1016/j.tree.2021.11.008.

Gabriel, L., Hoff, K. J., Brůna, T., Borodovsky, M., and Stanke, M. (2021). TSEBRA: transcript selector for BRAKER. BMC Bioinformatics 22, 566. doi: 10.1186/s12859-021-04482-0.

Gofas, S., Moreno D., Salas, C. (2012). Moluscos marinos de Andalucia - vol. 1.

Grossu, A.V. (1993). The catalogue of the molluscs from Romania. Trav. Mus. Hist. nat. Grigore Antipa, 33, 291 –366.

Hawkins, S. J., Mieszkowska, N., and Mrowicki, R. (2023). The genome sequence of the common limpet, Patella vulgata (Linnaeus, 1758). Wellcome Open Res. 8, 418. doi: 10.12688/wellcomeopenres.20008.1.

Hoff, K. J., Lange, S., Lomsadze, A., Borodovsky, M., and Stanke, M. (2016). BRAKER1: unsupervised RNA-Seq-based genome annotation with GeneMark-ET and AUGUSTUS. Bioinformatics 32, 767–769.

Hoff, K. J., Lomsadze, A., Borodovsky, M., and Stanke, M. (2019). “Whole-Genome Annotation with BRAKER,” in Gene Prediction: Methods and Protocols Methods in Molecular Biology., ed. M. Kollmar (New York, NY: Springer), 65–95. doi: 10.1007/978-1-4939-9173-0_5.

Hotaling, S., Kelley, J. L., and Frandsen, P. B. (2021). Toward a genome sequence for every animal: Where are we now? Proc. Natl. Acad. Sci. 118, e2109019118. doi: 10.1073/pnas.2109019118.

Karlicki, M., Antonowicz, S., and Karnkowska, A. (2021). Tiara: deep learning-based classification system for eukaryotic sequences. Bioinformatics 38, 344–350. doi: 10.1093/bioinformatics/btab672.

Kim, D., Paggi, J. M., Park, C., Bennett, C., and Salzberg, S. L. (2019). Graph-based genome alignment and genotyping with HISAT2 and HISAT-genotype. Nat. Biotechnol. 37, 907–915. doi: 10.1038/s41587-019-0201-4.

Koufopanou, V., Reid, D. G., Ridgway, S. A., and Thomas, R. H. (1999). A Molecular Phylogeny of the Patellid Limpets (Gastropoda: Patellidae) and Its Implications for the Origins of Their Antitropical Distribution. Mol. Phylogenet. Evol. 11, 138–156. doi: 10.1006/mpev.1998.0557.

Kuznetsov, D., Tegenfeldt, F., Manni, M., Seppey, M., Berkeley, M., Kriventseva, E. V., et al. (2023). OrthoDB v11: annotation of orthologs in the widest sampling of organismal diversity. Nucleic Acids Res. 51, D445–D451. doi: 10.1093/nar/gkac998.

Lawniczak, M. K., of Life, W. S. I. T., and Consortium, D. T. of L. (2022). The genome sequence of the blue-rayed limpet, Patella pellucida Linnaeus, 1758. Wellcome Open Res. 7. Available at: https://www.ncbi.nlm.nih.gov/pmc/articles/PMC9975397/ [Accessed December 8, 2023].

Lewin, H. A., Richards, S., Lieberman Aiden, E., Allende, M. L., Archibald, J. M., Bálint, M., et al. (2022). The Earth BioGenome Project 2020: Starting the clock. Proc. Natl. Acad. Sci. 119, e2115635118. doi: 10.1073/pnas.2115635118.

Lewin, H. A., Robinson, G. E., Kress, W. J., Baker, W. J., Coddington, J., Crandall, K. A., et al. (2018). Earth BioGenome Project: Sequencing life for the future of life. Proc. Natl. Acad. Sci. 115, 4325–4333. doi: 10.1073/pnas.1720115115.

Li, H. (2018). Minimap2: pairwise alignment for nucleotide sequences. Bioinformatics 34, 3094–3100. doi: 10.1093/bioinformatics/bty191.

Manni, M., Berkeley, M. R., Seppey, M., Simão, F. A., and Zdobnov, E. M. (2021a). BUSCO Update: Novel and Streamlined Workflows along with Broader and Deeper Phylogenetic Coverage for Scoring of Eukaryotic, Prokaryotic, and Viral Genomes. Mol. Biol. Evol. 38, 4647–4654. doi: 10.1093/molbev/msab199.

Manni, M., Berkeley, M. R., Seppey, M., and Zdobnov, E. M. (2021b). BUSCO: Assessing Genomic Data Quality and Beyond. Curr. Protoc. 1, e323. doi: 10.1002/cpz1.323.

Mazzoni, C. J., Ciofi, C., and Waterhouse, R. M. (2023). Biodiversity: an atlas of European reference genomes. Nature 619, 252–252. doi: 10.1038/d41586-023-02229-w.

Mc Cartney, A. M., Formenti, G., Mouton, A., Panis, D. D., Marins, L. S., Leitão, H. G., et al. (2023). The European Reference Genome Atlas: piloting a decentralised approach to equitable biodiversity genomics. 2023.09.25.559365. doi: 10.1101/2023.09.25.559365.

Menge, B. A. (2000). Top-down and bottom-up community regulation in marine rocky intertidal habitats. J. Exp. Mar. Biol. Ecol. 250, 257–289.

MolluscaBase eds. (2023). MolluscaBase. Patella caerulea Linnaeus, 1758. Available at: https://www.marinespecies.org/aphia.php?p=taxdetails&id=140677 [Accessed December 8, 2023].

Petraccioli, A., Guarino, F. M., Maio, N., and Odierna, G. (2010). Molecular cytogenetic study of three common Mediterranean limpets, Patella caerulea, P. rustica and P. ulyssiponensis (Archaeogastropoda, Mollusca). Genetica 138, 219–225. doi: 10.1007/s10709-009-9412-9.

Qi, W., Lim, Y.-W., Patrignani, A., Schläpfer, P., Bratus-Neuenschwander, A., Grüter, S., et al. (2022). The haplotype-resolved chromosome pairs of a heterozygous diploid African cassava cultivar reveal novel pan-genome and allele-specific transcriptome features. GigaScience 11, giac028. doi: 10.1093/gigascience/giac028.

Ranallo-Benavidez, T. R., Jaron, K. S., and Schatz, M. C. (2020). GenomeScope 2.0 and Smudgeplot for reference-free profiling of polyploid genomes. Nat. Commun. 11, 1432. doi: 10.1038/s41467-020-14998-3.

Reguera, P., Couceiro, L., and Fernández, N. (2018). A review of the empirical literature on the use of limpets Patella spp.(Mollusca: Gastropoda) as bioindicators of environmental quality. Ecotoxicol. Environ. Saf. 148, 593–600.

Rhie, A., McCarthy, S. A., Fedrigo, O., Damas, J., Formenti, G., Koren, S., et al. (2021). Towards complete and error-free genome assemblies of all vertebrate species. Nature 592, 737–746. doi: 10.1038/s41586-021-03451-0.

Rhie, A., Walenz, B. P., Koren, S., and Phillippy, A. M. (2020). Merqury: reference-free quality, completeness, and phasing assessment for genome assemblies. Genome Biol. 21, 245. doi: 10.1186/s13059-020-02134-9.

Ridgway, S. A., Reid, D. G., Taylor, J. D., Branch, G. M., and Hodgson, A. N. (1998). A cladistic phylogeny of the family Patellidae (Mollusca: Gastropoda). Philos. Trans. R. Soc. Lond. B. Biol. Sci. 353, 1645–1671. doi: 10.1098/rstb.1998.0316.

Sá-Pinto, A., Baird, S. J., Pinho, C., Alexandrino, P., and Branco, M. (2010). A three-way contact zone between forms of Patella rustica (Mollusca: Patellidae) in the central Mediterranean Sea. Biol. J. Linn. Soc. 100, 154–169.

Sá-Pinto, A., Branco, M., Harris, D. J., and Alexandrino, P. (2005). Phylogeny and phylogeography of the genus Patella based on mitochondrial DNA sequence data. J. Exp. Mar. Biol. Ecol. 325, 95–110.

Smit, A., Hubley, R., and Green, P. (2013). RepeatMasker Open-4.0. Available at: http://www.repeatmasker.org.

Storer, J., Hubley, R., Rosen, J., Wheeler, T. J., and Smit, A. F. (2021). The Dfam community resource of transposable element families, sequence models, and genome annotations. Mob. DNA 12, 2. doi: 10.1186/s13100-020-00230-y.

Theissinger, K., Fernandes, C., Formenti, G., Bista, I., Berg, P. R., Bleidorn, C., et al. (2023). How genomics can help biodiversity conservation. Trends Genet. 39, 545–559. doi: 10.1016/j.tig.2023.01.005.

Titselaar, F. F. (2019). Notes on the nomenclature of the Macaronesian Patella candei d’Orbigny complex, with special reference to Patella ordinaria Mabille and Patella crenata Gmelin (Patellogastropoda, Patellidae). Basteria 83, 158–165.

Vafidis, D., Drosou, I., Dimitriou, K., and Klaoudatos, D. (2020). Population characteristics of the Limpet Patella caerulea (Linnaeus, 1758) in Eastern Mediterranean (Central Greece). Water 12, 1186.

Zaidi, M., Athmouni, K., Metais, I., Ayadi, H., and Leignel, V. (2022). The Mediterranean limpet Patella caerulea (Gastropoda, Mollusca) to assess marine ecotoxicological risk: a case study of Tunisian coasts contaminated by metals. Environ. Sci. Pollut. Res. 29, 28339–28358. doi: 10.1007/s11356-021-18490-3.

